# Landmark knowledge overrides optic flow in honeybee waggle dance distance estimation

**DOI:** 10.1101/2024.06.10.598244

**Authors:** Randolf Menzel, C. Giovanni Galizia

## Abstract

Honeybees encode in their waggle dances the vector (distance and direction) of an outbound flight to a food source or a new nest side. Optic flow has been identified as the major source of information in the distance estimation. Additional components of distance estimation were also identified, e.g. the sequence of experienced landmarks. Here we address the question of whether bees also use the landscape memory developed during exploratory orientation flights to estimate distance. We took advantage of the fact that flights in a narrow tunnel lead to further distance measures due to higher optic flow. We find that this effect is lost when bees had explored the area in which the tunnel is located and when they have somewhat restricted visual access to the surrounding environment through the mesh on top of the tunnel. These data are interpreted in the context of other findings about the structure of navigational memory in bees that develops during exploratory orientation flights. In particular, the data suggest that bees embed distance measures into a representation of navigational space that stores previously experienced landscape features.

**Summary statement:** The distance code in the honeybee waggle dance is embedded in the landscape memory that bees establish during their exploratory and their foraging flights.

## Introduction

The waggle dance of honeybees encodes the distance and direction of the flight from the hive to a food source or to a new nest site (von Frisch, 1967). A most important discovery about the symbolic encoding of distance is the finding that the odometer of bees relies on optic flow during the outbound flight (Esch and Burns, 1995; Esch and Burns, 1996; Srinivasan et al., 2000). This was discovered by training bees in narrow tunnels that create higher optic flow than what is experienced during flight in the open environment. Increased optic flow leads to higher values of the distance code. In most experiments the distance measured by the bee was determined by the duration of the waggle run during the bees’ dances within the hive.

Additional and supportive information comes from experiments in which a feeder was located inside the tunnel and bees were videotaped during their search flight when the feeder was removed (Srinivasan et al., 1997). Combining these experimental approaches, it was possible to exclude alternative measures of flown distance, e.g. energy consumption (Heran, 1956; Heran, 1963), duration of flight, measuring and integrating airspeed, or some yet unknown measure of wing movement. Accumulating all this rich and supporting evidence (review: (Srinivasan, 2011)) it appears to be a well-established conclusion that the bee’s odometer receives its information only or predominantly from optic flow.

However, several observations indicate that additional or even alternative processes may also contribute to distance estimation. (1) Bees trained along serially placed landmarks fly to both the real distance of the feeder and the serially correct location if the distances between the landmarks were either increased or decreased (Chittka and Geiger, 1995; Menzel et al., 2010). Similar “counting “ effects of serially arranged marks were found when these marks were shown inside a tunnel (Dacke and Srinivasan, 2008a), indicating that both outside and inside the tunnel landmarks provide additional refence points for distance estimation. (2) Arranging a 6 m long tunnel in the open field at an angle of 90° to the direction of the approach flight did not lead to an accompanying shift of the danced waggle direction, rejecting the possibility that the flight in the tunnel contributes to a global vector based on a path integration process only (De Marco and Menzel, 2005). (3) Interestingly, such a global vector resulting from path integration was demonstrated by performing tunnel experiments in which the bees flew in the first half-length under a transversely oriented polarization filter (simulating a solar position that was directly ahead or behind the direction of flight), and the second half-length under an axially oriented polarization filter (simulating a solar position that was 90 deg to the left or the right of the flight direction) (Evangelista et al., 2014). These bees signaled a food source direction of 45 deg in their waggle dances, indicating an L-shaped flight with equal arm length, and thus integration of two paths under 90° direction. The waggle run duration of around 230 ms was found to be within the range of the results of (Srinivasan et al., 2000). Other than in the experiments of De Marco et al. (De Marco et al., 2008), the bees performed their outbound and inbound flights inside the tunnel and had most likely no access to external landmarks. (4) Srinivasan et al. (Srinivasan et al., 1997) found in experiments with the tunnel that a landmark inside the tunnel enhances the accuracy with which the bees searched for food, thus leading to a reduction of the error accumulation process in optic flow measures. (5) It is known that feeders closer to the hive are more attractive than more distant feeders of similar quality (e.g. sucrose concentration). Shafir et al. (Shafir and Barron, 2010) arranged two tunnels such that one was shorter than the other tunnel but induced higher optic flow (and thus should appear longer). Bees qualified the shorter tunnel better in their dances although it was associated with higher optic flow. (6) Dacke and Srinivasan (Dacke and Srinivasan, 2008b) concluded from their data that bees appear to have two odometers, one that drives waggle dance communication and one they use to estimate the total distance in their flights to a feeder they had visited before.

In all of these studies bees were trained to fly to a feeder in such a way that additional parameters besides optic flow competed with the distance estimation. Here, we have taken a different approach: we ask how the information from optic flow is integrated into what bees have learned during their previous exploratory flights at the beginning of their lives as foragers. Exploration of the environment is essential for bees before they start foraging (Capaldi et al., 2000; Degen et al., 2015). Sequential learning flights increasing to distances > 100 m and varying in direction lead to a knowledge of the environment surrounding the hive that allows them to find home from anywhere within the explored area via direct flights (Degen et al., 2016). The memory established during exploration is best understood as integrating egocentric, allocentric and compass information including local as well as global guiding cues (Menzel, 2023). Such a memory would potentially allow extracting a flown distance from this highly integrated form of spatial memory.

One may ask, therefore, how these different reference systems for distance estimation interact and under which conditions one dominates the other or whether compromises are made when information is inconsistent and bees have to communicate distance in the waggle dance. We address this question by setting up tunnel experiments under conditions in which the bees were differently familiar with the terrain in which the experiments were performed. For most of the experiments the colonies were positioned in the environment more than 4 weeks before the experiments started, ensuring that the foragers tested had explored the environment outside of the tunnel. The surroundings were characterized by rich landmarks (trees, bushes, houses). In one experiment they had explored a different environment and were relocated just before the experiment. We found that the familiarity with the environment resulting from exploratory flights (and possibly additionally from foraging flights to natural food sources), rather than optic flow information, dominated the distance communicated in the waggle dance.

## Materials and methods

The experiments were performed with observation hives (containing approximately 3,000 bees each) in the summers 2022 and 2023. An IR camera (Raspberry Pi) monitored the dance area close to the entrance/exit of the hive. The experimental site was a highly structured domestic area in the village Amöneburg (Germany, 50°47’35.7”N 8°55’36.9”E) with trees, bushes, houses, roads.

The flight tunnels were rather similar to those used by Srinivasan et al. (Srinivasan et al., 2000), with length varying between 0.5 m and 6 m (in the preliminary experiment) and 6 m (in the main experiments 1 to 7) with an inner width of 11 cm, and a height of 30 cm. The top of the tunnel was covered with a metal-colored insect-screen (Fig. 1). Bees saw the sky above them and rising landmarks in the surrounding within an angle approx. up to 60° during their flights in the tunnel. In the main experiment bees saw the surrounding environment only during the flight in the tunnel and not at all when they were feeding at the feeder F, because a light tight box was mounted at the end of the tunnel containing the feeder allowing to observe and monitor the marked bees (Fig. 1). The floor and the sides of the tunnel were covered with a black and white random texture with pixel size of 1 cm by 1 cm. Bees were trained to a feeder located outside, at the entrance of the tunnel or the end of the tunnel depending on experimental design (Fig. 2, 3).

**Fig. 1.**
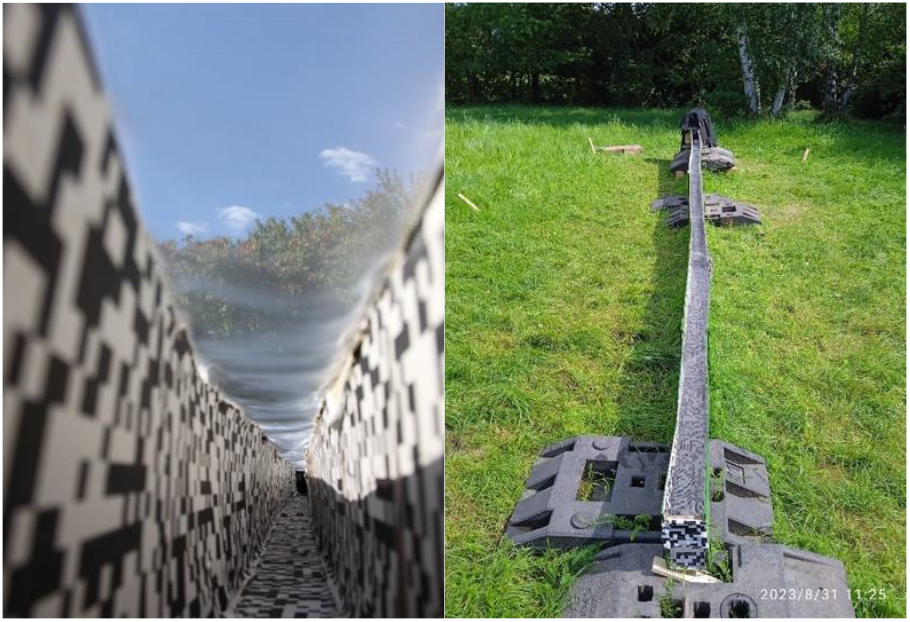
Left: view from within the tunnel as used during the preliminary experiments and the experiments 1 – 6 of the main experiment. Right: view of the tunnel from outside. Note the concealed ending: the feeder was in the dark, and the experimenter could enter the cover.

**Fig. 2.**
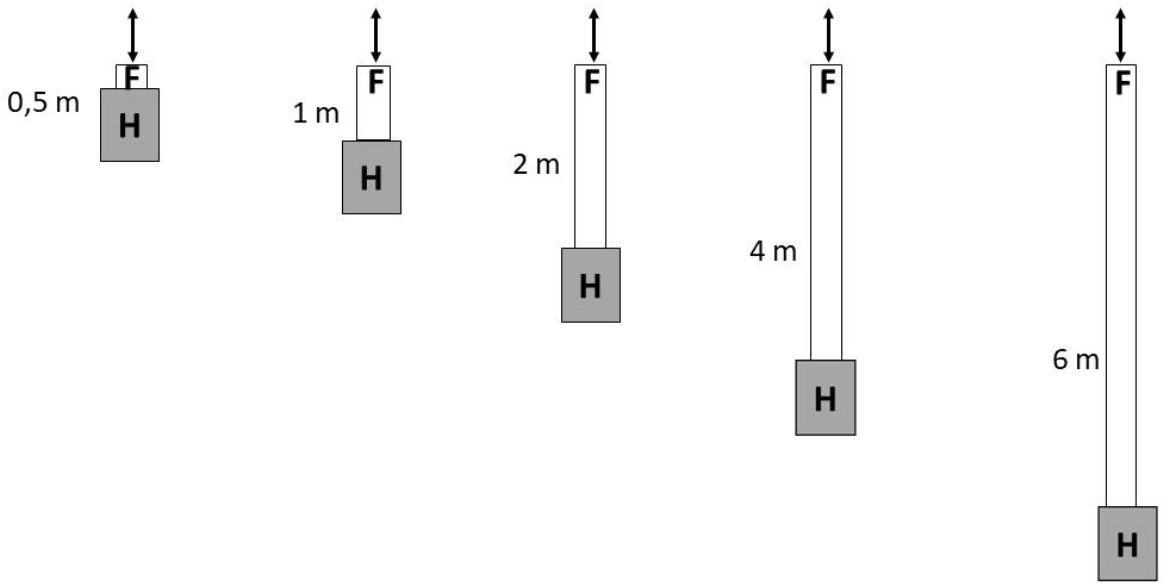
Preliminary experiments. The colony in the observation hive had long time experience with the environment. Tunnels of different length (0.5, 1, 2, 4 or 6 m) were attached to the front of the hive with the entrance hole such that the entrance/exit to the tunnel was always at the same location. The feeder was close to the end inside the tunnel. The length of the tunnel was changed several times by moving the colony accordingly. Bees visiting the feeder were marked with a white dot at the abdomen. A monitor displaying the images of the IR video camera recording the bees’ dances within the hive was set up behind the hive, and dances of the marked bees were visually observed and evaluated. Bees not trained to the feeder (thus not marked) were free to move in and out at the end of the tunnel and were not included in the on-line evaluated dances.

**Fig. 3.**
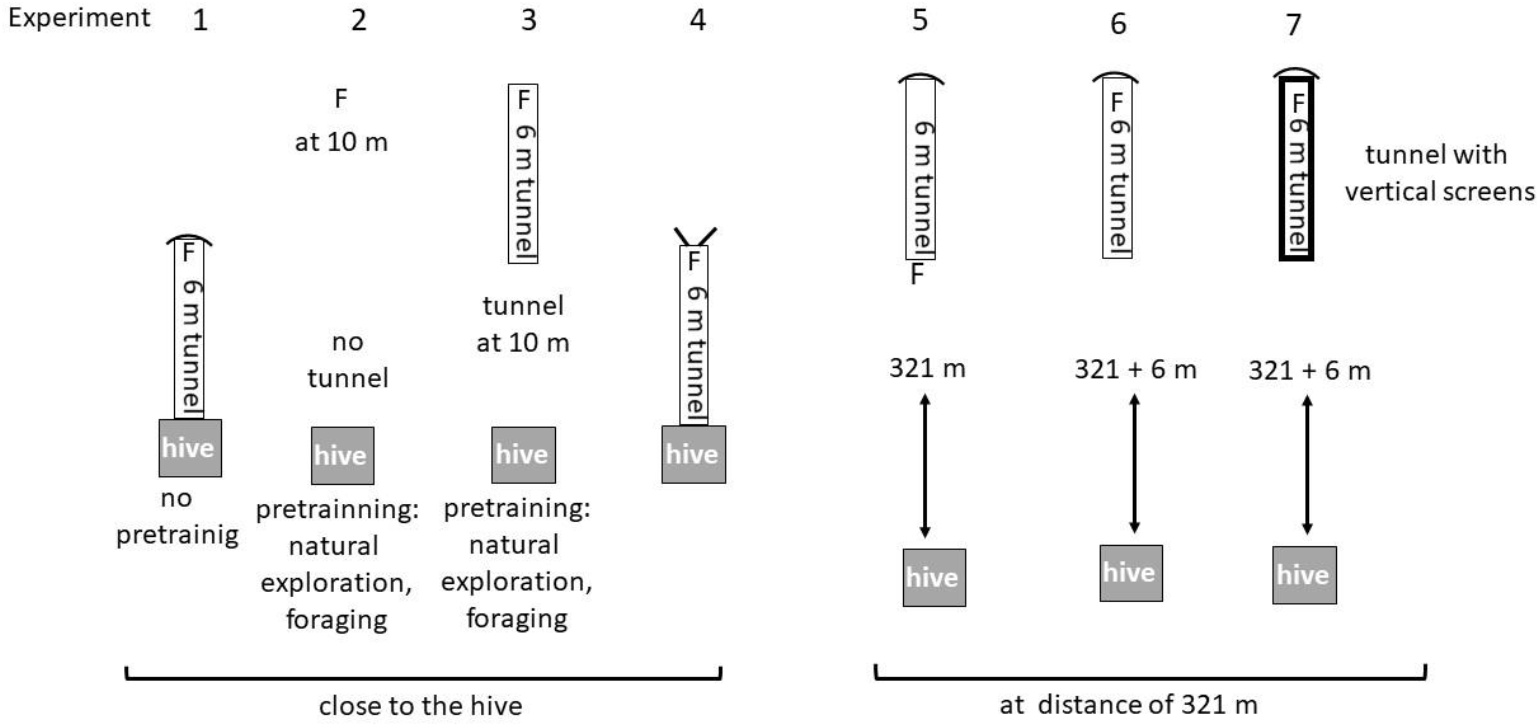
Design of the experiments in the main experiment. Experiment 1: The tunnel was attached to the entrance of the hive prior to any exploratory flights, foraging flights or feeder (F) training. Experiment 2: The foraging bees were trained to a feeder at a distance of 10 m from the hive. Experiment 3: Experienced foragers were trained to the end of the 6 m tunnel. The entrance to the tunnel was at 10 m from the hive. Experiment 4: The tunnel was attached to the entrance of the hive after the bees had explored the environment and foraged at natural food sources. Experiment 5: experienced foragers were trained to a feeder (F) at a distance of 321 m and subsequently fed at this location. Experiment 6: bees from experiment 5 were trained to a feeder (F) at the end of the 6 m tunnel. Experiment 7: The animals from experiment 5 and 6 were further trained to the feeder of the tunnel but two screens (2.5 m high) were attached to the side walls of the feeder. In this situation the animals could see the sky but no landmarks surrounding the feeder. in experiment 4 indicates that the entrance/exit to the hive via the tunnel is open for other foragers not feeding at F, marks the closed end of the tunnel that is covered with a box allowing access to the feeder and blocking the view of the surrounding during feeding at F.

Two sets of experiments were run. In the first set (preliminary experiments, Fig. 2) the tunnels were of different length (0.5, 1, 2, 4 or 6 m) and attached to the entrance/exit of the hive such that the end of the tunnel was always at the same location relative to the external landmarks. The length of the tunnel was changed several times by moving the colony accordingly. A feeder was always located at the end of the tunnel. The far end of the tunnel was open to allow foraging bees not taking part in the experiment to fly in and out freely. Bees visiting the feeder were marked with a white dot at the abdomen. Within the hive, dances of the marked bees were visually observed via a video camera. A monitor displaying the images of these recordings was set up behind the hive. The colony in the observation hive had long time experience (at least 4 weeks) with their environment.

In the main experiments the location of the tunnel, the location of the feeder and the far end of the tunnel (open or closed) varied according to the individual design of the different experiments (experiments 1 – 7, Fig. 3). Bees visiting the feeder were individually marked with dual digit black and white number tags (or only pre-marked for experiment 1). The number range for differently marked bees was enhanced to 470 by positioning the tags in 4 different directions on the thorax. The dances on the dance floor were video-taped using an IR Raspberry Pi camera. The videos were analyzed off-line with the help of a custom written video analysis script in Python that detected the location and the time of a waggle run, stopped the video and opened a window that allowed to mark the start and the end of the waggle run as detected by the first and the last frame in which the bee was unsharp due to her fast-waggling movement. The video was recorded with 50 frames per second, and bees appeared somewhat unsharp during the waggle run but not during normal walking or return runs, allowing to set the frames for start and end of the waggle run accurately. The video frame was calibrated for space and time and the following data were noted in the pop-up window of the program and saved to file: duration of the waggle run, its length and the number of waggles performed, and the direction of the waggle run relative to gravity. The latter was used by the script to derive the angle to north in reference to the location of the hive, the date and the time of the day. This procedure led to efficient and precise measurements of large numbers of waggle runs. Furthermore, these data allowed to compare the variance and the correlation of two possible codes of distance: the duration and the number of waggles per waggle run. We found that number of waggles per waggle run varied less than waggle duration, and therefore used this metric (see results section, Fig. 4). The correlation between number of waggles and duration of the waggle run allowed us to relate our data to published data that used duration as the distance code.

**Fig. 4.**
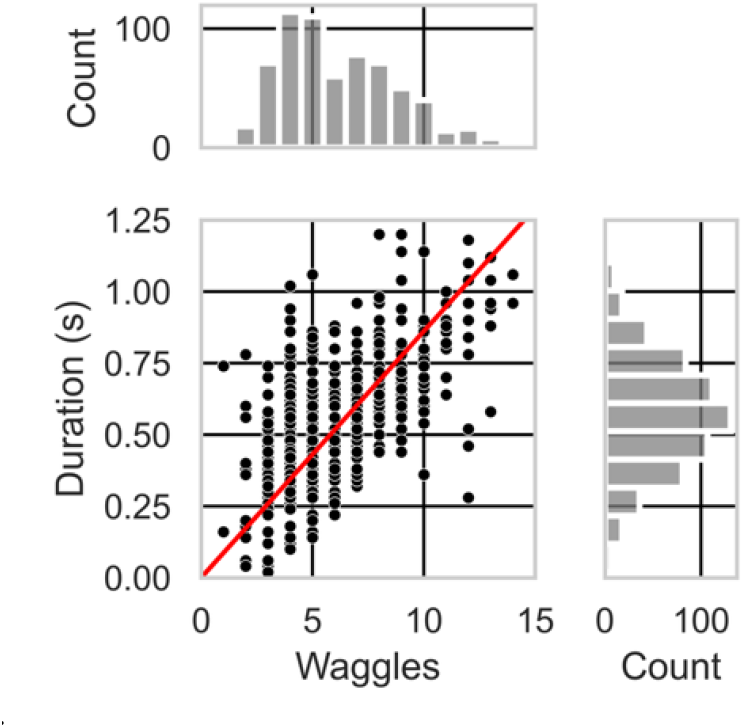
Plot of waggles/run against duration/run, for experiments 1, 6 and 7 of the main experiment group, with histograms for both at the side. Note that the histogram for waggles has two peaks: at 5 waggles/run corresponding to experiments 1 (n=84, mean±std: 4.1±1.1) and 6 (n=233, mean±std: 4.3±1.3), and at 7 waggles/run corresponding to experiment 7 (n=337, mean±std: 8.1±2.1). The higher variability in duration smears the distribution to a single peak (mean±std for duration, Exp. 1, 6, 7: 0.43±0.18, 0.48±0.19, 0.67±0.18).

The tunnels of the main experiment were close to the hive in the experiments 1 – 4 and at a further distance (321 m, coordinates: 50°47’46.1”N 8°55’35.5”E) in the experiments 5 – 7. As mentioned above the bees flying in the tunnel could see the external environment within an angle of approximately 60° during their flights in the tunnel because they always flew close underneath the mesh. They did not see the environment during feeding. In experiment 1, the colony was located first in an area about 4.5 kilometers from the experimental area behind a hill (50°48’52.3”N 8°52’20.7”E). Many foraging bees were marked with a white dot at the abdomen before the colony was moved. Then they were moved over night into the experimental area, the 6 m long tunnel was attached to the hive and the far end closed. Thus, the colony was naïve to the environment because it was moved into the test environment shortly before the experiment started and the bees had not explored the test area yet. The feeder could be inserted and refilled without allowing bees to fly out or to approach it from the outside.

Thus, foragers feeding at the end of the tunnel in experiment 1 had only experience with the tunnel and not with the environment around the tunnel. This was different in experiments 2 to 7. The foragers in these experiments had explored the environment. They could reach the feeder only by flying through the tunnel. Experiments 2 and 5 were control experiments with a feeder at 10 m distance from the hive (experiment 2) or at 321 m distance (experiment 5) and no tunnel flight. In experiment 3 the entrance of the 6 m tunnel was at a distance of 10 m and the feeder was located at the end of the tunnel. In experiment 4 the tunnel was attached to the hive. The difference to experiment 1 was that the bees had experience with the surroundings before flying through the tunnel to the feeder at the end of the tunnel. In experiment 6, bees visited the feeder at the end of the tunnel. In experiment 7, two screens (2.5 m high) tightly attached to the right and left of the tunnel excluded the view of landmarks outside the tunnel but left the view to the sky.

Statistics and plotting were done using *Python* 3.9.15, *Pandas* 2.1.4, *Seaborn* 0.13.2, *Statsmodel* 0.14.0, *SciPy* 1.11.3. Boxplots show quartiles, whiskers the full distribution, except for outliers that are determined using a method in *Seaborn* that is a function of the inter-quartile range.

## Results

### Number of waggles per waggle run as a code for distance

Our initial objective was to compare various parameters of the waggle run to identify which of them had the least variation, and thus which would deliver the most accurate distance code (recent review (Kohl and Rutschmann, 2021)). The number of dance rounds in 15 seconds, commonly utilized by Karl von Frisch (von Frisch, 1967) in many experiments, exhibited high variability; the length of the waggle run varied also considerably (data not shown). Consequently, both measures were excluded from our analysis. Instead, we concentrated on two parameters for the same waggle runs: the number of waggles per run and the duration of a run. Both parameters were measured for the same waggle runs through video analysis in a subset of our data, as described above. The duration of the waggle run displayed slightly more variability than the number of waggles per run (p<0.01, Levene Test). It is noteworthy that the frequency distribution of the durations was close to a single Gaussian distribution and did not show any indication of a double peaked distributions (Fig. 4, right histogram), while the frequency distribution of waggle count/run showed two distinct peaks, one for experiments 1 and 6, and another one for experiment 7 (Fig. 4, top histogram), revealing that the lower variability in this metric kept the two distinct distributions visible. Therefore, for the bulk of our measurements, we evaluated the waggle count metric. A linear regression gave a slope of 0.086 s per waggle.

### Dances during the preliminary experiments

The design of the preliminary experiment allowed to distinguish between round dances and waggle dances. The observation hive was positioned on a trolley, allowing for quick mobility of the hive while the entrance/exit of the tunnel and the feeder at the end of the tunnel remained stationary. This is a necessary requirement because bees learn the surroundings of the hive very accurately, and in these experiments all bees accessed the hive via the tunnel irrespective of its length and whether they were trained to a feeder at the end of the tunnel or flew to natural food sources. Outbound and inbound bees not feeding at the feeder accommodated very quickly to the changing length of the tunnel. Thus, the entrance/exit to the observation hive via the tunnel remained stable, while the length of the tunnel varied (see Fig. 2). These test conditions facilitated the alteration of tunnel length, ensuring that bees encountered the access to the hive via the tunnel while maintaining rather constant spatial relations to the environment. Based on the literature we expected waggle runs under 4 of the 5 test conditions (i.e. in all conditions with the tunnels longer than 0.5 m).

Prior to the start of the preliminary experiments, foragers from the colony experienced a condition with a short tunnel (0.5 m) for several weeks. A feeder was positioned 10 cm from the entrance/exit within the tunnel, and bees visiting the feeder were identified by marking them with a white dot on the abdomen. The tunnel’s length was modified once at least 50 dances were observed. Multiple rounds of semi-random insertions of tunnels with varying lengths were conducted, and the feeder was always 10 cm from the entrance/exit of the tunnel irrespective of its length. Only round dances, and no waggle dances, were observed in marked bees for all 5 test conditions. This is an important and rather surprising finding because one would have expected that under the conditions of this experiment either only waggle dances were performed in four of the five test conditions, or an increasing number of waggle runs with increasing tunnel lengths. Thus, our results falsified the hypothesis we had in mind when we started this experiment (see above).

A substantial difference to the experiments of (Srinivasan et al., 2000) was that our tunnel flying bees could see the surrounding environment which they had learned before during their exploratory orientation flights. Furthermore, although the bees feeding at the feeder inside the tunnel and close to its far end predominantly shuttled between the feeder and the access to the hive at the other end of the tunnel, some of them may have flown out of the tunnel from time to time since the far end of the tunnel was not closed. These considerations lead to designing the main experiment in which the potential effects of the exploratory experience with the natural environment prior to the tunnel flights were systematically tested.

### Dances during experiments 1 – 7 in the main experiment

Seven experiments were run in the main experiment (Fig. 3). In experiment 1, the colony in the observation hive was first located for 5 weeks in an area approximately 4.5 km away from the experimental area (50°48’52.3”N 8°52’20.7”E). The landscape here (agricultural fields, grass land) was very different from that of the experimental area (domestic area in the village). In the last week before moving the hive, many foragers were marked with a white dot on the abdomen. The forgers were not trained to a feeder, and the natural food supply was scattered over larger distances (> 200 m) and rather scarce. Before the experiment, the hive was moved during the night, a 6 m tunnel was attached to the entrance/exit, and the tunnel was closed at the far end. A feeder was placed at the far end within the tunnel, and a small shelter allowed to examine the feeder and refill it. Videos of the dance floor were recorded with an IR camera and the dances analyzed off-line using the procedure described in the Method section. In experiment 2 – 7 the colony in the observation hive was located in the experimental area for 3 weeks before the experiment started.

In experiment 1 three sessions of 30 minutes recording each were analyzed. The marked bees predominantly performed waggles dances (n=298, waggles/run mean±std: 4.1±1.1, Fig. 5); 2 round dances were observed. Thus, flights in the 6 m tunnel with view to an unexplored environment led to waggle runs, indicating that the optic flow in the tunnel elicited a long-distance dance (see below for calibration).

**Fig. 5.**
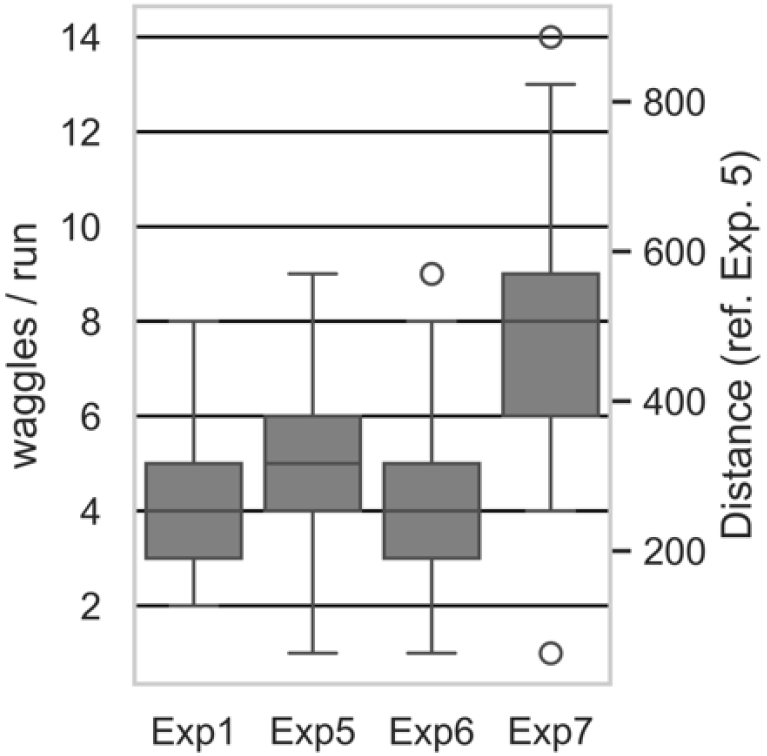
Distribution of waggles/run in experiments 1, 5, 6 and 7. Boxes show quartiles, whiskers rest of distributions, except for outliers (see methods). Right ordinate axis shows distances in meters indicated by the waggle dance, calibrated with experiment 5 (no tunnel). (see S2 for data and statistics).

No tunnel was used in experiment 2, and the feeder was located 10 m from the hive in the direction of 351° to N. Four sessions of 30 minutes each were video recorded. We observed only round dances but no waggle runs, in accordance to the short distance flown. However, the same result was found also for experiments 3 and 4: only round dances. In experiment 3 the entrance/exit to the 6 m tunnel was at distance of 10 m from the hive, direction 354° to N, feeder at the end of the tunnel, and in experiment 4, the 6 m tunnel was attached to the hive and the feeder at the end of the tunnel. This latter experiment required a different colony because a new colony had to be brought into the experimental area, and the foragers learned to access the hive through the tunnel. Therefore, experiment 4 was carried out at the time when colony 1 was exposed to a different environment (experiment 1). The results show that in experiments 3 and 4, bees danced for a location in the immediate vicinity, despite of having flown through the tunnel (which simulated long-distance in experiment 1). Therefore, in the next experiments 5-7, we asked under which conditions does a tunnel simulate a long-distance, and when does it not.

In experiments 5 – 7, foragers familiar with the landscape were trained to a distant location (321 m, direction 354° to N). In experiment 5 the feeder was at the entrance of the 6 m tunnel (bees did not fly through the tunnel) and in experiment 6 at the end of the 6 m tunnel. In experiment 7 the feeder was also at the end of the tunnel but two screens (2.5 m high) excluded the view of landmarks outside the tunnel and left the view to the sky. As expected, foragers performed waggle dances in all these conditions (Fig. 5).

In experiment 5, bees danced about 5 waggles/run to indicate the 321 m distance (n=227, mean±std: 5.1±1.5); in experiment 6, with the added 6 m tunnel, the waggles did not increase (n=547, mean±std: 4.2±1.2); in experiment 7, when shielding the 6 m tunnel from the surroundings, the waggles increased (n=456, mean±std: 7.6±2.2; Fig. 5). A generalized linear model analysis (Poisson model family, log link function, IRLS method with post-hoc testing) showed no significant difference between the results in experiment 1 and 6.

Experiment 5 and experiment 7 differed significantly (p < 0.001). Even though the difference between experiments 5 against 1 and 6 is significantly different, the ranges are strongly overlapping (see Fig. 5), suggesting that this difference may not have a biological relevance. However, the distribution ranges of experiment 7 are clearly distinct (Fig. 5): here flying through the tunnel let to a highly relevant and significant increased distance as signaled in the waggle dance.

We used the data from experiment 5 to calibrate the distance code for number of waggles: each waggle/run indicated a 63.3 m distance (approx. 5 waggles/run for the known 321 m distance, see Fig.5). Applying this calibration, bees indicated a fictive distance for the 6 m tunnel of 262 m in experiment 1. A similar distance was danced in experiment 6 (265 m, short of the really flown distance of 321+6 m). In experiment 7, bees danced 481 m. If we subtract the open distance of 321 m, this indicates a danced tunnel length of 220 m, i.e., slightly shorter than in exp. 1 (see right ordinate scale in Fig. 5). This experiment also shows that bees add up distance in free flight with the distance within the tunnel, when flying both sequentially. Taken together, these data suggest that the environment surrounding the tunnel provides information for distance coding if the dancer is familiar with the environment (in experiment 6 the bees did not experience the tunnel as a long distance, while in experiment 7, with no view of the environment, the tunnel was experienced as a long distance).

| Exp. 1 |  | Exp. 5 | Exp. 6 | Exp. 7 |
| --- | --- | --- | --- | --- |
| 4.1 ± 1.1 | waggles | 5.1 ± 1.5 | 4.2 ± 1.2 | 7.6 ± 2.2 |
| 298 | n | 227 | 547 | 456 |
| 0.356 s | duration (calc) | 0.437 s | 0.360 s | 0.655 s |
|  | m/waggle | 63.3 m | 78.1 m |  |
|  | danced distance (ref exp 5) | 321 m | 265 m | 481 m |
| 262 m | danced tunnel length (5) | 59 m | 3 m | 220 m |
|  | danced distance (ref exp 6) | 396 m | 327 m | 594 m |
| 323 m | danced tunnel length (6) | 73 m | 4 m | 271 m |

The observations in the preliminary experiments (Fig. 3) and the video analyses of experiments 2, 3, 4 and 6 showed clearly that the input from the environment dominated the distance coding when the dancers had explored the environment. However, it could be that the average values of waggles per run in these experiments may have resulted from some sort of switching between the competing inputs. We plotted the distribution of waggles/run for each experiment in order to investigate whether there was any evidence for independent dual information (Fig. 7). We found that the frequency distribution was close to Gaussian with no indication of double peaked distributions in any of the experiments.

Marking individual bees with number tags allowed us to further address the question whether individual dancers exposed to competing conditions may differ in coping with this situation. We had marked 368 foragers with number tags and hoped that they would forage in both experiments 6 and 7 allowing us to see at the individual level whether they would deal with the test conditions differently (Fig. 7). Unfortunately, no tagged dancers were seen in our videos that were exposed to both test conditions. However, calculating the average of waggles per waggle run for each of the individuals separately for the two test conditions allowed us to reject the possibility that some individuals may have weighted the two inputs differently.

## Discussion

The flight through the 6 m tunnel simulated a flight distance of 262 m (experiment 1, tunnel close to the hive) or a distance of 220 m (experiment 7, tunnel further away from the hive) if environmental information at the test site was excluded, thus confirming the finding that optic flow is a major factor for bees to estimate flight distance. The data reported in (Srinivasan et al., 2000) (their Fig. 2, experiments 2 and 4) indicated optic-flow induced distances of a similar 6 m tunnel of 184 m (close to the hive) and 230 m (further away from the hive). These results are rather close to the result reported here given the condition that different colonies were used and the tunnels had different heights. Our tunnel was 30 cm high and their tunnel 20 cm high. The corresponding durations of waggle runs in (Srinivasan et al., 2000) were 441 ms for the tunnel close to the hive and 529 ms for the tunnel further away from the hive, which is in the same order as the results found here: applying the conversion of number of waggles to duration of 0.086 s per waggle (Fig. 4) we obtained 356 ms waggle run duration for the close tunnel (Exp. 1) and 408 ms for experiment 7 (Fig. 6).

**Fig. 6.**
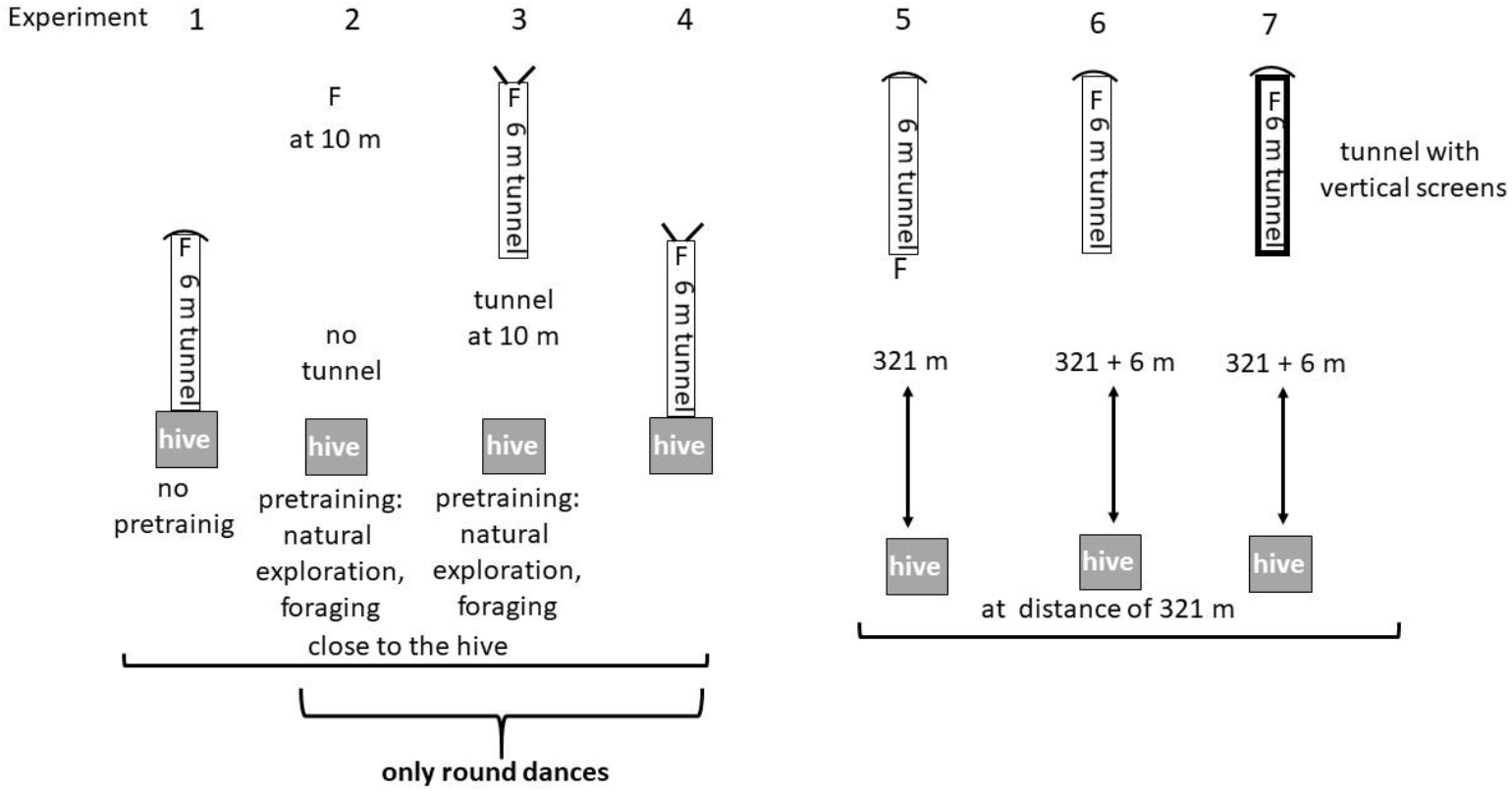
Summary of the results on dances in the main experiment. The upper part of the figure repeats the design of the 7 experiments, the lower part gives the data for the experiments 1, 5, 6 and 7 together with the calculations of distances from the number of waggles and the corresponding durations as calculated from Fig. 4.

**Fig. 7.**
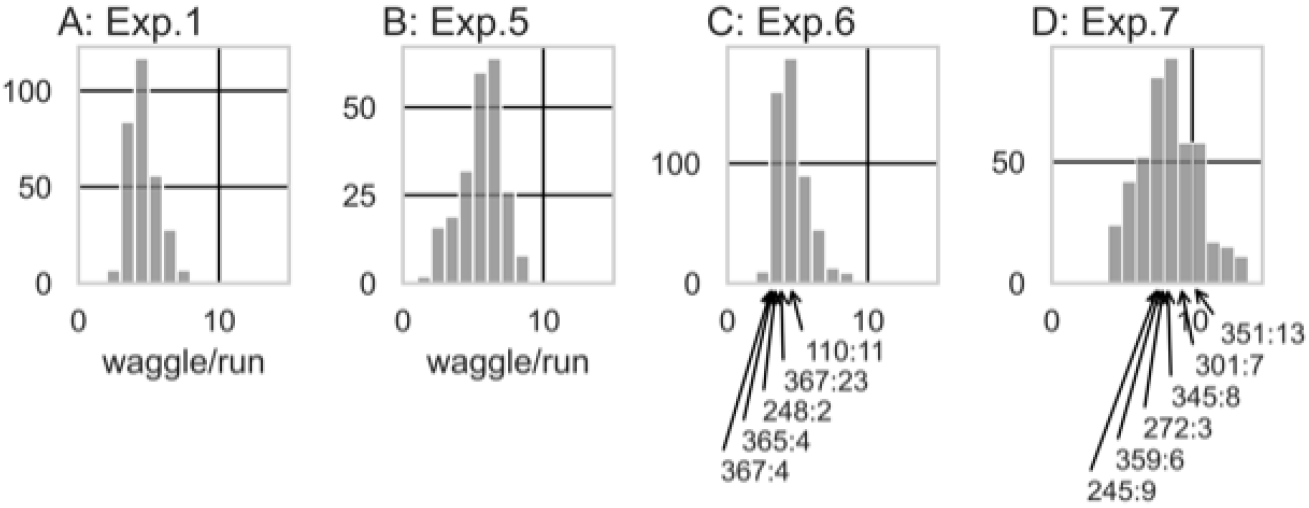
Histograms of waggles/run in experiments 1, 5, 6 and 7. In each experiment, a single prominent peak is visible. Note the high values in experiment 7 as compared to experiments 1, 5 and 6. In experiments 6 and 7, the values for identified bees are indicated with arrows toward the abscissa, the label indicates the bee identity, and the number of averaged (observed) waggle dances (e.g.: bee number 110 had 11 evaluated dances in experiment 6).

The data presented here add an important component to distance estimation in honeybees that goes beyond the measurement and the encoding/decoding of distance in the waggle dance. We show that knowledge of the environment surrounding the tunnel can override the optic flow effect. This partly corroborates findings cited in the introduction with respect of effects of serial landmark learning (Chittka et al., 1995; Menzel et al., 2010), “counting” phenomena (Dacke and Srinivasan, 2008a), reduced error accumulation through serial landmarks (Srinivasan et al., 1997), and importantly the discovery of two odometers in bees (Dacke and Srinivasan, 2008b). In addition, the study by (De Marco and Menzel, 2005) showed that large range path integration collapses when bees fly out of the tunnel that was arranged 90° to the access flight to the feeder. Interestingly, unlike the results presented here, the optic flow effect was not overridden in that study when the bees continued flying the same direction inside of the tunnel as in the access flight, even though the view of the environment was not blocked. This can be explained by the special conditions of the environment around the tunnel in their study. The experiments were carried out in a large, flat and horizontal grassland without rising objects and a flat horizon. The bees saw the environment only in the moment when they left the tunnel, and in that moment their knowledge of the environment took over. The differences between (Evangelista et al., 2014) and (De Marco and Menzel, 2005) on path integration can also be resolved on the basis of our data reported here. Flights only inside the tunnel with little or no view of the environment as in the case of the Evangelista 2014 (Evangelista et al., 2014) study restrict the distance measure to optic flow, whereas the moment the bees leave the tunnel and return back to the hive in flight through the open they will refer their distance measure to the landscape memory.

What we have termed “knowledge of the environment” and “landscape memory” here should be understood as a technical term catching the consequences of exploratory learning. Conceptually, this memory could have two forms within the bees’ neural networks: either, each point of the known environment is elementally associated to a homing vector, or, different points of the known environment are connected in a navigational map. The elemental hypothesis would mean that in our experiments bees referred to a list of independent memories storing associations between single and specified landscape features associated with the homing vector. Such an elementary form of landscape memory cannot strictly be excluded by these experiments. However, this might be quite costly for the bee brain in terms of necessary memory capacity, and offers little flexibility in natural environments with all their daily and seasonal changes. We argue, therefore, that the data reported here can be explained in a more parsimonious way, i.e. using a navigational map. Exploratory learning of a navigational map differs from elemental target associative learning in several important aspects (Birke and Archer, 1983; Gallistel, 1990; Renner, 1988; Tolman, 1948). The process of exploration is an attention inducing and rewarding process in itself accompanied with active movement. Sequentially experienced and spatially separated objects are bound together leading to a representation of organized space, and multiple experiences of similar cues (both of the egocentric and allocentric domain) will make the spatial memory richer and more precise (Chen and Mou, 2024; Hilton and Wiener, 2023). It has been argued that multiple exploratory flights lead to memory storage and retrieval processes that appear to bind together separate memories through generalization process, memory updating, completion and correction (Menzel, 2023). Taken together these multiple and independent data sets allow to interpret the memory structure resulting from exploration as being substantially different from memory resulting from associative learning. In our view, this form of spatial/temporal representation is best captured by the term navigational map, sometimes referred to as cognitive map (Jeffery et al., 2024; Tolman, 1948). Extraction of elementary components from such global representations, e.g. the distance between objects in explored space, requires the retrieval of specified memory of the appropriate part of space and its spatial connection to other locations, e.g. the center of life (nest) or recently visited locations (feeding places). The accumulating supporting evidence in favor of such a form of spatial representation in waggle dance followers allows to conclude that the endpoint of the symbolically encoded flight vector (distance and direction) is represented as a location in this spatial memory (Wang et al., 2023). The kind of cognitive operations are performed most likely both in waggle dancers and in waggle dance followers because dancers frequently switch between foraging, dance following and dancing. Taken together, we conclude that the measure of distance as expressed in the dance is embedded in the global representation of the explored space. Phenomena like dancing for a food source after a detour flight (e.g. around a mountain (von Frisch, 1967), p. 174 – 178) or uphill could mean that dancers and followers estimate the true distance (further distance) by referring their flight to the learned characteristics of the landscape. In an ecological context, trips need to be planned taking into account changing properties of the environment and weather conditions.

## Acknowledgements

We are grateful to Manu Dür for his help with the Python script that helped to analyze the videos of dancing bees, and Dr. Jana Mach for her help in programming of the Raspberry Pi cameras.

## Data availability

All data will be placed in a public repository upon publication.

## References

Birke, L. I. and Archer, J. (1983). Some issues and problems in the study of animal exploration. Exploration in animals and humans, 1–21.

Capaldi, E. A., Smith, A. D., Osborne, J. L., Fahrbach, S. E., Farris, S. M., Reynolds, D. R., Edwards, A. S., Martin, A., Robinson, G. E., Poppy, G. M. et al. (2000). Ontogeny of orientation flight in the honeybee revealed by harmonic radar. Nature 403, 537–540.

Chen, Y. and Mou, W. (2024). Path integration, rather than being suppressed, is used to update spatial views in familiar environments with constantly available landmarks. Cognition 242, 105662.

Chittka, L. and Geiger, K. (1995). Can honeybees count landmarks? Anim. Behav 49, 159–164.

Chittka, L., Geiger, K. and Kunze, J. (1995). The influences of landmarks on distance estimation of honey bees. Anim. Behav 50, 23–31.

Dacke, M. and Srinivasan, M. V. (2008a). Evidence for counting in insects Anim Cogn 11, 683–689.

Dacke, M. and Srinivasan, M. V. (2008b). Two odometers in honeybees? J Exp. Biol 211, 3281–3286.

De Marco, R. J., Gurevitz, J. M. and Menzel, R. (2008). Variability in the encoding of spatial information by dancing bees J Exp. Biol 211, 1635–1644.

De Marco, R. J. and Menzel, R. (2005). Encoding spatial information in the waggle dance. J. Exp. Biology 208, 3885–3894.

Degen, J., Kirbach, A., Reiter, L., Lehmann, K., Norton, P., Storms, M., Koblofsky, M., Winter, S., Georgieva, P. B., Nguyen, H. et al. (2016). Honeybees Learn Landscape Features during Exploratory Orientation Flights. Current Biology 26, 2800–2804.

Degen, J., Kirbach, A., Reiter, L., Lehmann, K., Norton, P., Storms, M., Koblofsky, M., Winter, S., Georgieva, P. B., Nguyen, H. et al. (2015). Exploratory behaviour of honeybees during orientation flights. Animal Behaviour 102, 45–57.

Esch, H. E. and Burns, J. E. (1995). Honeybees use optic flow to measure the distance of a food source. Naturwiss 82, 38–40.

Esch, H. E. and Burns, J. E. (1996). Distance estimation by foraging honeybees. Journal of Experimental Biology 199, 155–162.

Evangelista, C., Kraft, P., Dacke, M., Labhart, T. and Srinivasan, M. (2014). Honeybee navigation: critically examining the role of the polarization compass. Philosophical Transactions of the Royal Society B: Biological Sciences 369, 20130037.

Gallistel, C. R. (1990). The Organization of Learning. Cambridge, Mass., London: MIT Press.

Heran, H. (1956). Ein Beitrag zur Frage nach der Wahrnehmungsgrundlage der Entfernungsweisung der Bienen. Z. vergl. Physiol 38, 168–218.

Heran, H. (1963). Wie beeinflu?t eine zusätzliche Last die Fluggeschwindigkeit der Honigbiene? Verh. Dt. Zool. Ges. Wien 26, 346–354.

Hilton, C. and Wiener, J. (2023). Route sequence knowledge supports the formation of cognitive maps. Hippocampus 33, 1161–1170.

Jeffery, K. J., Cheng, K., Newcombe, N. S., Bingman, V. P. and Menzel, R. (2024). Unpacking the navigation toolbox: insights from comparative cognition. Proceedings of the Royal Society B 291, 20231304.

Kohl, P. L. and Rutschmann, B. (2021). Honey bees communicate distance via non-linear waggle duration functions. PeerJ 9, e11187.

Menzel, R. (2023). Navigation and dance communication in honeybees: a cognitive perspective. Journal of Comparative Physiology A, 1–13.

Menzel, R., Fuchs, J., Nadler, L., Weiss, B., Kumbischinski, N., Adebiyi, D., Hartfil, S. and Greggers, U. (2010). Dominance of the odometer over serial landmark learning in honeybee navigation Naturwissenschaften 97, 763–767.

Renner, M. J. (1988). Learning During Exploration: The Role of Behavioral Topography During Exploration in Determining Subsequent Adaptive Behavior in the Sprague-Dawley Rat (Rattus norvegicus). International Journal of Comparative Psychology 2.

Shafir, S. and Barron, A. B. (2010). Optic flow informs distance but not profitability for honeybees. Proceedings of the Royal Society B: Biological Sciences 277, 1241–1245.

Srinivasan, M. V. (2011). Honeybees as a model for the study of visually guided flight, navigation, and biologically inspired robotics. Physiological reviews 91, 413–460.

Srinivasan, M. V., Zhang, S., Altwein, M. and Tautz, J. (2000). Honeybee Navigation: Nature and Calibration of the” Odometer". Science 287, 851–853.

Srinivasan, M. V., Zhang, S. W. and Bidwell, N. J. (1997). Visually mediated odometry in honeybees. Journal of Experimental Biology 200, 2513–2522.

Tolman, E. C. (1948). Cognitive maps in rats and men. Psychol. Rev 55, 189–208.

von Frisch, K. (1967). The dance language and orientation of bees. Cambridge: Harvard Univ.Press.

Wang, Z., Chen, X., Becker, F., Greggers, U., Walter, S., Werner, M., Gallistel, C. R. and Menzel, R. (2023). Honey bees infer source location from the dances of returning foragers. Proceedings of the National Academy of Sciences 120, e2213068120.

